# The structure of the iron-catecholate transporter Fiu suggests substrate import occurs via a 2-step mechanism

**DOI:** 10.1101/763268

**Authors:** Rhys Grinter, Trevor Lithgow

## Abstract

The Ferric Iron Uptake (Fiu) transporter from *Escherichia coli* functions in the transport of iron-catecholate complexes across the bacterial outer membrane, providing the bacterium with iron which is an essential element for growth. Recently, it became clear that Fiu also represents a liability: its activity allows the import of antimicrobial compounds that have evolved to mimic catecholate. In this work we have determined the structure of Fiu and analyzed its function to address how Fiu and related transporters from other bacterial species can bind catecholate in a surface-exposed cavity. In addition, the crystal structure of Fiu reveals the presence of a large, selectively gated cavity in the interior of this transporter. This chamber is large enough to accommodate the Fiu substrate and may act to regulate substrate import. These data provide insight into the mechanism of substrate uptake by Fiu and related transporters identified in *Pseudomonas aeruginosa* and *Acinetobacter baumannii*. As Fiu and its homologues are the targets of substrate mimicking antibiotics, these data will assist in the development of antibiotics that target these receptors for cell entry.

## Introduction

The outer membrane of Gram-negative bacteria provides a selective permeability barrier to molecules with a molecular mass greater than ∼600 Da [1]. The barrier provides superb protection against antimicrobials and toxic compounds [2]. The selective permeability of the outer-membrane also restricts the uptake of nutrients including iron which, while abundant on Earth, is often growth-limiting due to its insolubility under the oxidizing conditions of the terrestrial atmosphere [3, 4]. Aerobic organisms solubilize iron through formation of iron-chelating chemicals (siderophores) or incorporate it into other organic structures such as the porphyrin ring of heme or iron-binding proteins [5–7]. These iron-containing complexes are larger than the diffusion limit of the bacterial outer-membrane, and so in order to obtain the iron required for growth, bacteria have evolved transporters capable of selectively binding and importing of iron-containing complexes [8, 9]. Members of the TonB-dependent transporter (TBDT) family drive transport of their substrates through interaction with the energy transducing protein TonB [10]. TBDTs are highly divergent in sequence, yet share a common structural architecture consisting of a 22-stranded transmembrane β-barrel, the lumen of which is selectively occluded by a globular plug domain [8].

The ability of TBDTs to import large substrates comes at a cost: an evolutionary arms race exists between TBDT-producing bacteria and organisms seeking to kill them by hijacking these receptors [11–15]. Both small molecule and protein antibiotics mimic TBDT substrates, leading to their inadvertent import into the bacterial cell [12, 16–21]. Catecholates are one of the four recognized classes of siderophores [22], have a strong affinity for iron, and are abundant secretion products of both bacteria and fungi [23, 24]. The Ferric Iron Uptake (Fiu) transporter is an TBDT responsible for the import of molecules containing the catecholate functional group [2, 25]. In addition to its role in iron uptake, Fiu has also been shown to be important for sensitivity to antimicrobials that share the common feature of a catecholate functional group or the analogous dihydroxypyridine moiety [21, 25, 26]. It is thought that the presence of these chemical mimics leads to their inadvertent import into the bacterial cell [27]. This Fiu mediated sensitivity is observed even in the absence of iron, because Fiu functions in the import of catecholate containing molecules seemingly independent of their size or Fe-coordination state [18]. The potential of antimicrobial catechol-siderophore mimetics as therapeutics is demonstrated by the development of the 3,4-dihydroxypyridine containing sulfactam BAL30072 by Basilea Pharmaceutica, and the catecholate to containing Cefiderocol cephalosporin by Shionogi Inc. BAL30072 and Cefiderocol have entered clinical trials for the treatment of infections by Gram-negative bacteria, in the human lung and urinary tract respectively [28, 29].

In this study, we show that while Fiu and its homologues PiuA and PiuD function in the import of catechol-siderophores [25, 30], they are evolutionarily distinct from the well-studied catecholate siderophore transporters FepA/PfeA and Cir. In order to investigate the substrate import mechanism of the Fiu/Piu TBDT subgroup, we solved the crystal structure of Fiu. Analysis of this structure in combination with *in silico* docking and mutagenesis identified an external substrate binding site in Fiu, which is conserved among diverse TBDTs. In addition, the presence of a large selectively gated internal chamber in Fiu, capable of accommodating an Fe-siderophore complex, suggests these transporters may function via an airlock style gating mechanism.

## Results

### Fiu is a member of a distinct clade of iron-catecholate transporters

It was previously demonstrated that the archetypical *E. coli* strain BW25113 possesses three TBDT transporters that function in the uptake of catecholate siderophores: FepA, Cir and Fiu [25]. FepA imports the endogenously produced siderophore enterobactin with high affinity [31], while Cir and Fiu have been shown to transport monomeric catecholate compounds, either alone or in complex with iron [32]. While these transporters recognize a common functional group, they share limited amino acid sequence identity and their evolutionary relationship remained undetermined. To resolve this question, we performed phylogenetic analysis on these transporters in the context of a panel of diverse TBDTs of known structure and/or function. This analysis revealed that while Cir and FepA belong to the same clade of our TBDT phylogram, Fiu belongs to a distal clade with the TBDTs PiuA and PiuD that also mediate catecholate transport [30, 33] (Figure 1A). Wider analysis of this Fiu/Piu clade, identified by HMMER search reveals that related sequences are widespread in proteobacteria (Figure S1, Table S1, Table S2). Based on clustering analysis of these sequences, all members of this group are more closely related to Fiu than either Cir or FepA, and consistent with our phylogram are more closely related to the hydroxamate siderophore transporter FhuA (Figure 1A, Figure S1). These data demonstrate that while Fiu, Cir and FepA all transport catecholate containing substrates, Fiu is evolutionarily distinct from Cir and FepA, and may have arrived at its substrate specificity due to convergent evolution between these transporters.

**Figure 1.**
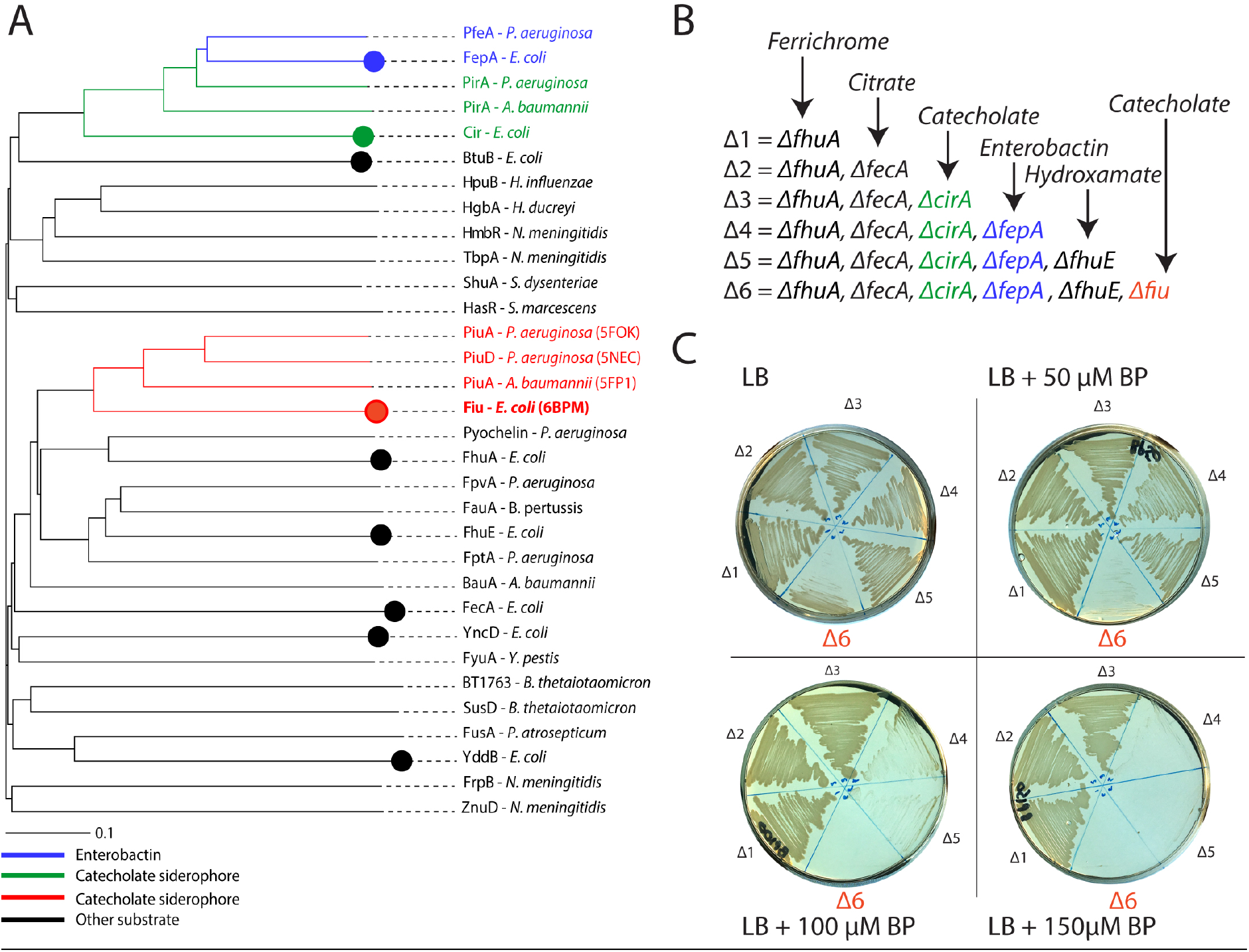
Fiu belongs to a distinct group of catecholate siderophore transporters. (A) A phylogenetic tree of diverse functionally characterised TBDTs, showing that Fiu forms a clade with PiuA/D, that is distant from the catecholate siderophore transporters FepA and Cir. (B) A scheme showing the sequential deletion TBDTs in *E. coli* BW25113 utilised in this study. (C) Strains from panel B, grown on LB agar in the presence of 0-150 μM 2’2-bipyridine (BP). Sequential loss of FepA and Fiu leads to defects in the ability of strains to grow under iron limiting conditions.

TBDT mediated iron-uptake systems are generally redundant in order to provide the means to obtain iron under a variety of environmental conditions [23, 34, 35]. Thus, to dissect the specific role of Fiu, each of the six TBDTs known to be involved in iron acquisition were sequentially deleted from *E. coli* BW25113 in the following order: Δ*fhuA* (ferrichrome transporter), Δ*fecA* (ferric-citrate transporter), Δ*cirA* (Fe-catecholate siderohpore transporter), Δ*fepA* (enterobactin transporter), Δ*fhuE* (rhodotorulic acid transporter) and finally Δ*fiu* (Figure 1B). The phenotypes of these mutants were assessed by growth on LB agar containing the iron chelator 2’2-bipyridine (Figure 1C). The first three receptors (FhuA, FecA and CirA) were dispensable for growth in this assay, but the subsequent loss of the enterobactin receptor FepA affected the growth of mutant strain at 2’2-bipyridine concentrations of > 50 μM (Figure 1C). There was no further phenotype from loss of the coprogen receptor FhuE under our assay conditions (Δ5; Figure 1C). Subsequent loss of Fiu, led to impaired growth on LB agar and completely prevented growth at 2’2-bipyridine concentrations 50 μM or higher (Δ6; Figure 1C). This growth defect was restored either by in-trans complementation with a plasmid encoding Fiu (Figure S2) or supplementation of the growth medium with Fe(II)SO_4_.

These data show that Fiu is able to provide iron to the cell when present as the sole outer membrane iron transporter. As Fiu is unable to transport endogenously produced enterobactin [31, 36], in this context it most likely functions to transport enterobactin breakdown products (i.e. 2,3-dihydroxybenzoyl-L-serine (DHBS)) in complex with iron. The inability of Fiu to support growth in the presence of high concentrations of 2’2-bipyridine, may be due to the lower affinity of the monomeric catecholates for Fe^3+^ or a low affinity of Fiu for the Fe-DHBS complex.

### Crystal structures of Fiu reveals a large, gated internal chamber

To obtain insight into the structural basis for substrate binding and import by Fiu, we determined the structure of Fiu by X-ray crystallography (Table S3). The structure of Fiu consists of a 22-stranded transmembrane β-barrel characteristic of the TBDT superfamily, with a number of extended extracellular loops that might serve in initial steps of substrate binding (Figure 2A). In agreement with our phylogenetic analysis (Figure 1A), the closest structural homologue of Fiu in the PDB was PiuA from *A. baumannii* (PDB ID = 5FP1, Z-score = 45 and RMSD = 2.0), which shares 33 % amino acid sequence identity with Fiu. The structure of Fiu was solved in three different crystal forms (Table S3), revealing Fiu in two distinct states. In crystal state 1 extracellular loops 7-9 of the β-barrel were disordered, as was the extended extracellular loop of the N-terminal plug domain, which occludes the lumen of the β-barrel. While in crystal state 2, amino acids 50-760 of Fiu could be traced into the electron density; beginning immediately C-terminal of the TonB box, which is disordered in our structure, and extending to the end of the Fiu polypeptide (Figure S3A). The disorder of the plug domain loop in crystal state 1, opens a large cavity in the interior of Fiu to the external environment, while in crystal state two this cavity is present, but occluded in the lumen of the Fiu barrel (Figure 2B). Interestingly, analysis of the structures of PiuA from *Acinetobacter baumannii* and, PiuA and PiuD from *P. aeruginosa* revealed analogous crystal states to Fiu (Figure S3A) [30, 33]. In PiuA from *A. baumannii* all extracellular loops are ordered with the plug loop occluding the entrance to an internal cavity (Figure S3). While in PiuA and PiuD from *P. aeruginosa* the extracellular and plug loops are disordered, with the external cavity of PiuA exposed to the external environment (Figure S3).

**Figure 2.**
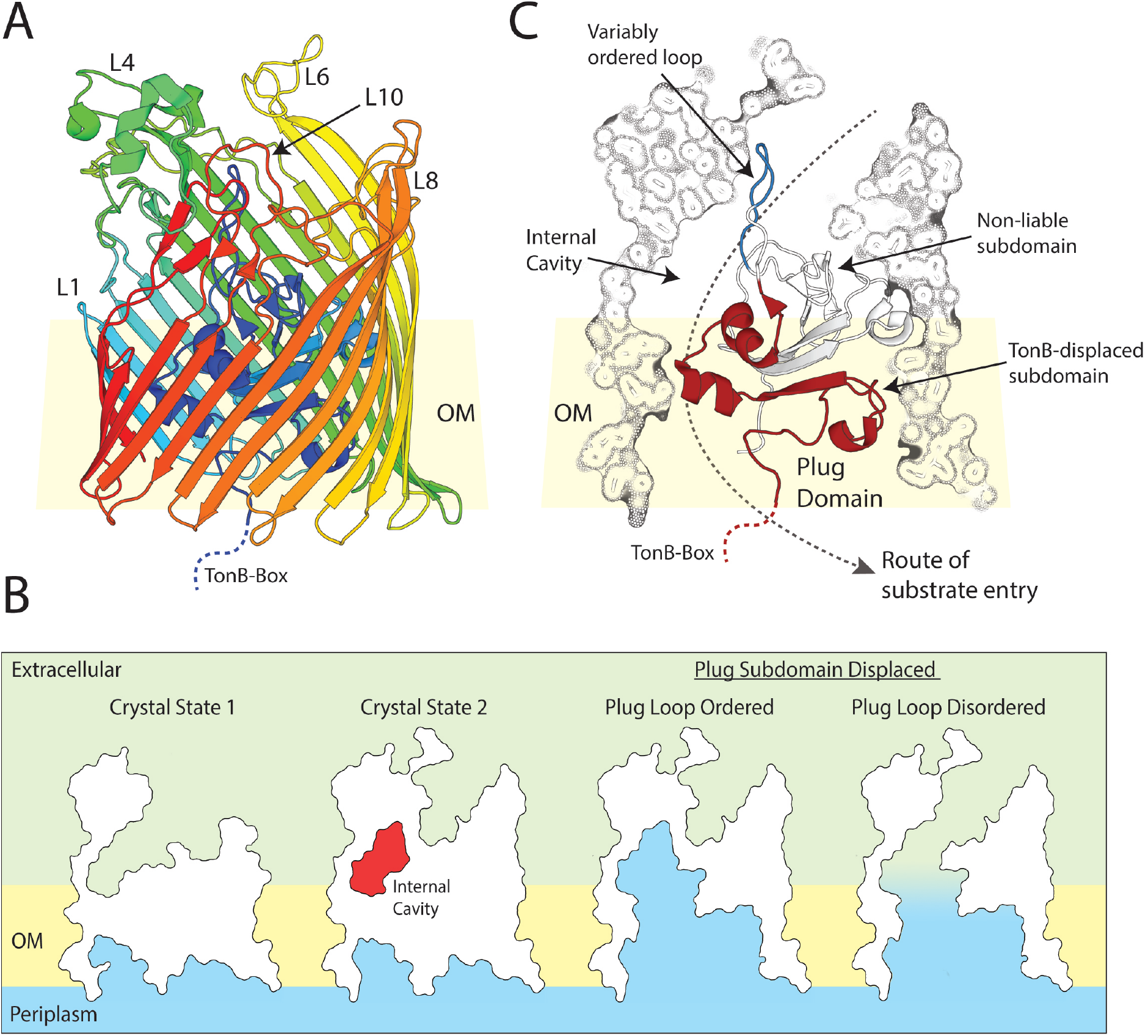
The crystal structure of Fiu reveals a large gated internal cavity. (A) The crystal structure of fully ordered Fiu (crystal state 2), shown as a cartoon representation with rainbow color running from N-terminal (blue) to C-terminal (red). (B) A cutaway outline representation of Fiu crystal structure showing the selectively occluded internal cavity in crystal states 1 and 2, as well as the result of the removal of the liable plug subdomain on Fiu channel formation through the membrane. (C) A composite cutaway view of Fiu showing the N-terminal plug domain as cartoon, with variably ordered plug loop (blue) and liable subdomain (red) highlighted.

Utilizing force spectroscopy Hickman et. al. showed that N-terminal plug domain of TBDTs consists of liable and non-liable subdomains [37]. Upon substrate binding, TonB is recruited to the TonB-box at the N-terminus of the TBDT and facilities the removal of liable subdomain, via mechanical energy provided the proton motif force [37, 38]. Prior to this study, a number of charged residues at the interface between the liable and non-liable subdomains in TBDT were shown to be important for substrate transport, but not binding [39]. Based on these studies, we identified that the liable subdomain of the Fiu plug extends from the N-terminus of the protein, to the start of extracellular plug domain loop that is selectively ordered in our crystal structures (Figure 2C). In crystal state two removal of the liable subdomain opens the internal cavity of Fiu to the periplasm, but due to the presence of the plug loop this does not create a membrane spanning channel. While in crystal state 1 removal of this subdomain opens a large channel between the periplasm and the external environment (Figure 3B).

**Figure 3.**
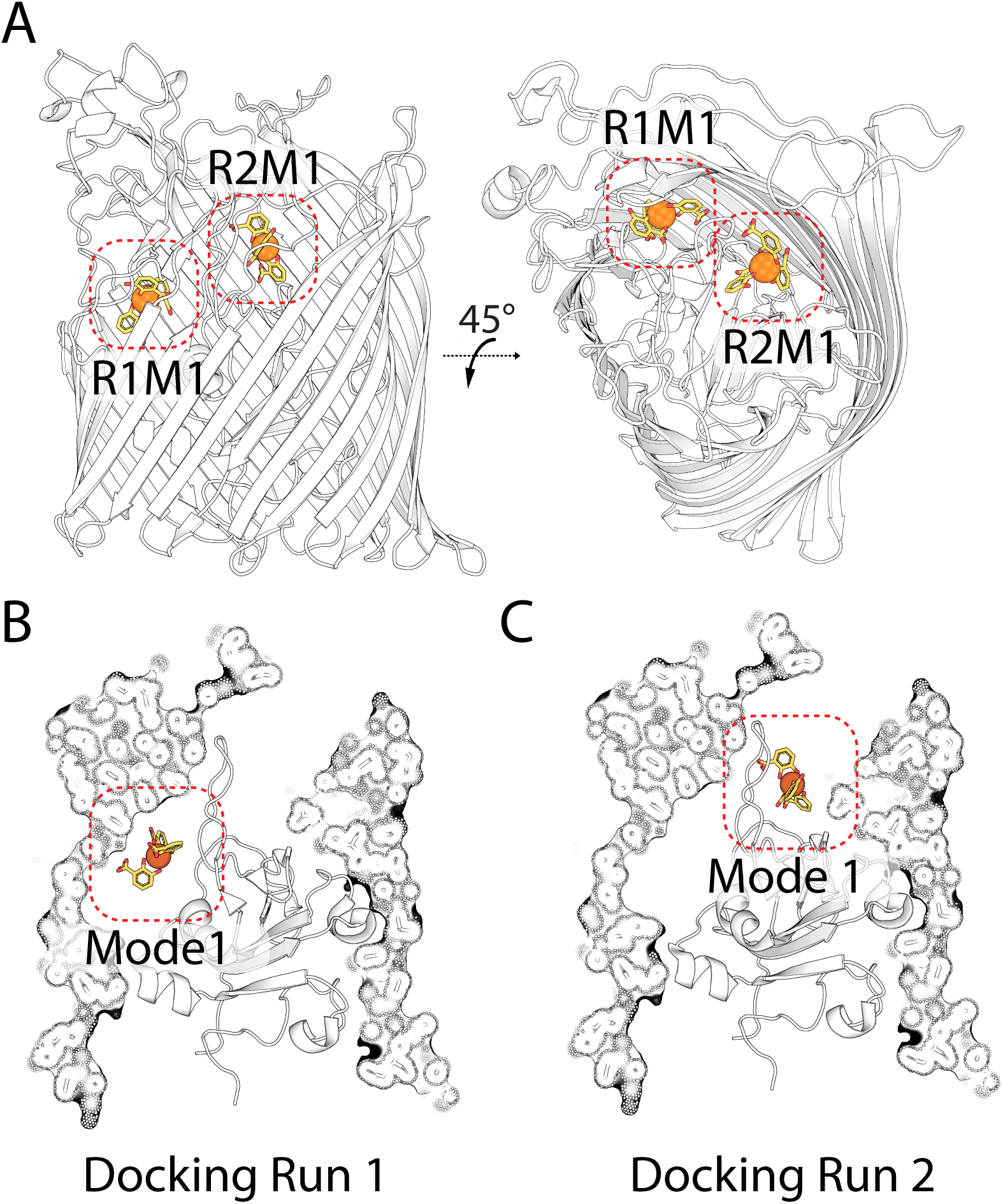
Top ranked docking solutions between Fiu and Fe-DHB substrate. (A) The location of the top ranked docking modes in a cartoon representation of Fiu, for docking run 1 (R1M1) including the entire extracellular portion of the receptor, and docking run 2 (R2M1) with the internal cavity excluded from the search area. (B) The same docking solutions as panel a shown on a cutaway composite view of Fiu.

TBDTs selectively transport their substrate across the outer membrane while, excluding antibiotics and other deleterious molecules from entering the cell [8]. As such, it is likely to be undesirable for Fiu to exist in the open channel state, which would result from simultaneous displacement of the plug subdomain and disorder of the plug domain loop. The internal cavity we observed in our structure, gated by the selectively ordered plug loop may provide a solution to this problem. The internal cavity is large enough (∼3200 Å^3^) to accommodate a Fe-siderophore complex, and so if siderophore entered this chamber prior to remove of the plug subdomain, it could enter the periplasm without the formation of a membrane spanning channel through the pore of Fiu.

### *In silico* docking suggests Fiu possesses multiple substrate binding sites

In order to determine the substrate binding site of Fiu, we attempted co-crystallization and soaking of Fiu crystals in the presence of the monomeric catecholate compounds DHBS and 2,3-dihydroxybenzoic acid (DHB) in complex with Fe^3+^. These monomeric catecholates form a 3:1 complex with a single Fe^3+^ ion at the center. Despite the presence of DHBS at a high concentration (100-1000 μM) during crystallization screening and soaking, and the resulting Fiu crystals diffracting well (2.8-2.0 Å), no electron density corresponding to DHBS was observed in the resulting density maps. Some crystals of Fiu grown in the presence of high concentrations of Fe-DHB (1 mM), exhibited the characteristic purple color of the Fe-catecholate complex (Figure S4A). While these crystals only diffracted to low resolution (anisotropic diffraction 3.2-5 Å), Fo-Fc density and anomalous difference density attributable to two Fe-DHB complexes was observed (Figure S4B). However, these Fe-DHB complexes were located on the side of the Fiu barrel, distal from the extracellular binding pocket and were involved in crystal packing, suggesting they are bound non-specifically (Figure S4C). While it has been demonstrated previously that Fiu is capable of transporting DHB and DHBS *in vivo* [32], our inability to obtain a legitimate co-crystal structure suggests that Fiu has a low affinity for these compounds. As TBDTs generally bind their ligands with very high affinity [8], this suggests that monomeric catecholate compounds may not be the preferred substrate for Fiu and its target siderophore remains unidentified.

To determine potential substrate binding sites in Fiu, we performed *in silico* docking between Fiu and Fe-DHB using Autodock Vina [40]. The rationale for this experiment is that while DHB appears to be a low affinity ligand, the ability of Fiu to transport it suggests that the high affinity substrate for this transporter is likely a catecholate siderophore, which would contain a Fe^3+^-catecholate complex analogous to DHB. Thus, while results should be interpreted cautiously, docking with Fe-DHB provides an indication substrate binding sites in Fiu. Two docking runs were performed, for the first run the entire extracellular portion of closed Fiu (crystal state 2) was defined as the search area. In this experiment, the majority of the solutions placed Fe-DHB inside the internal chamber of Fiu, with the 3^rd^ most favored solution positioning Fe-DHB in the extracellular cavity of the protein (Figure 3A/B, Figure S5, Table S4). While the internal cavity would be inaccessible to Fe-DHB in the closed state, due to the fully ordered plug loop, it would be accessible in state 1 as this loop is disordered. In the second docking experiment the internal cavity was excluded from the search area. In this experiment all solutions placed Fe-DHB in the Fiu extracellular cavity, with the majority of solutions clustered at a single location (Figure 3A/C, Figure S5). Suggestively, the top-ranking solution from this docking run was identical to 3^rd^ top solution from the first experiment. Taken together these results suggest that Fiu possesses a binding site capable of accommodating a Fe-catecholate complex in its extracellular cavity. In addition, these data show that the internal cavity of Fiu is capable of accommodating the Fe-DHB complex.

### The location of the putative external Fiu substrate binding site is conserved among TBDTs

In order to validate the external substrate binding site identified in our docking analysis, we compared its location to the substrates of other TBDTs that have been structurally characterized. For this analysis we selected a 12 non-redundant TBDTs-ligand structures (Table S5) [38, 41–49] and superimposed them with the structure of Fiu. In 11 of these 12 structures the substrate bound in an analogous location to our Fe-DHB docking solution, with the metal ion of the respective ligands located between 2.8 and 9.5 Å from the Fe in the docked complex (Figure 4). The 12^th^ structure, the ligand of which did not colocalize with Fe-DHB, is the enterobactin transporter PfeA from *P. aeruginosa* (Figure 4). In this structure the binding of enterobactin in PfeA occurs on the external face of the extracellular loops of the transporter, which entirely enclose the entrance to the lumen of the barrel [42]. It was demonstrated that PfeA binds enterobactin via 2-site binding and that the site observed in this crystal structure represents the initial substrate binding site, with the second binding site located deeper inside the transporter barrel [42]. This 2-site binding model is further supported by the structural analysis of FepA, a close homologue of PfeA, which shows that it binds enterobactin at two locations, the one of which is analogous to the PfeA binding site [50]. Fiu and the other TBDTs analyzed lack the extracellular loop structure required to bind their substrates in this external location, and so it is likely that the binding site observed in PfeA and FepA is distinct to other TBDTs. These data suggest that TBDTs share a common substrate binding site that is analogous to the Fiu external binding site identified by the docking analysis, providing validation of this result.

**Figure 4.**
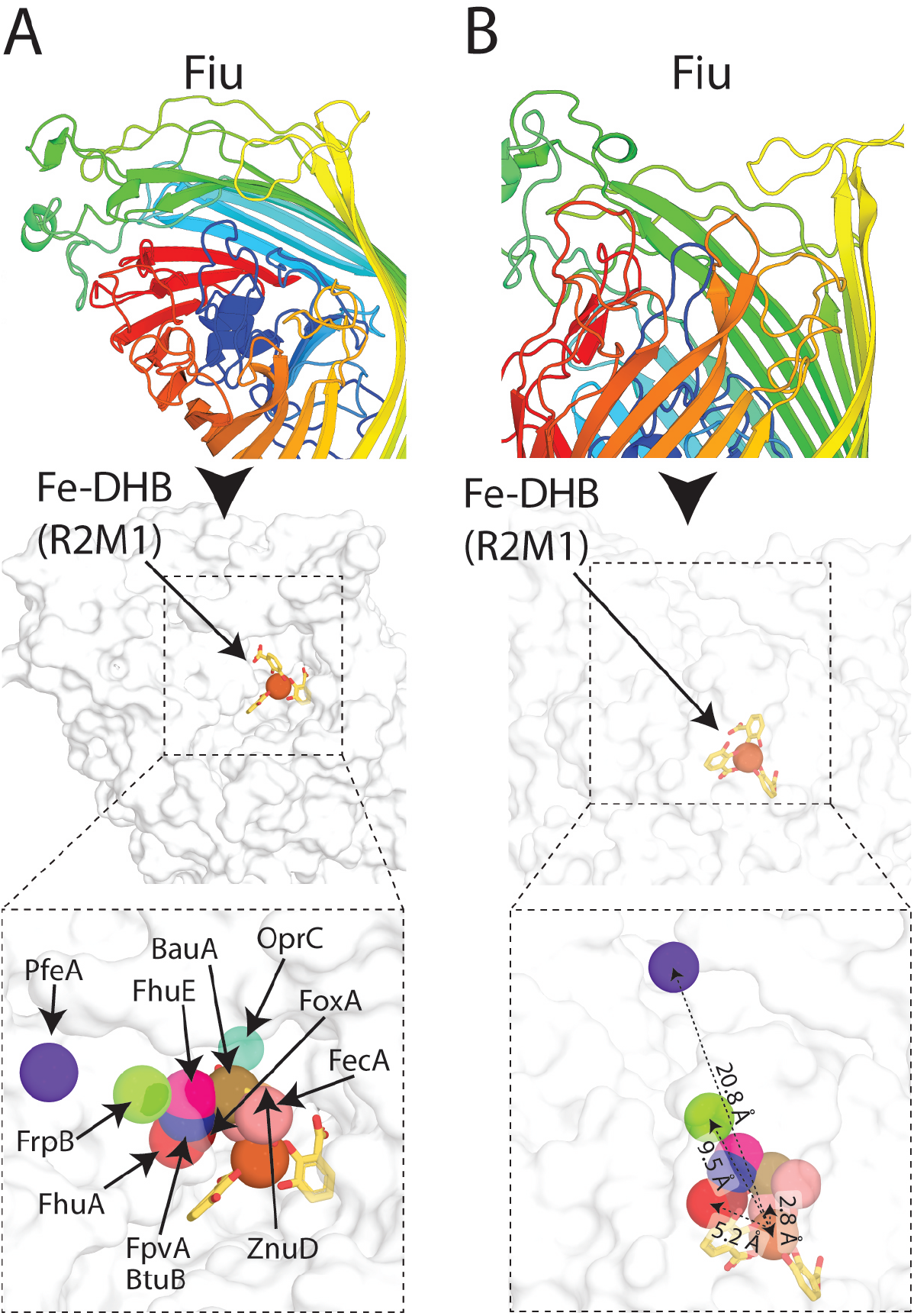
The location of the Fiu putative external Fe-DHB binding site compared to other TBDT substrate complexes. (A) The location of the Fiu Fe-DHB docked complex, in comparison to that of other substrates bound to in the crystal structures of other TBDTs superimposed with Fiu. Fiu is shown as a cartoon rainbow, and in the same view as a white surface representation below. The location of the metal centers of different TBDT substrates are shown as colored spheres and labeled. (B) The Fiu Fe-DHB docked complex shown as in panel A, at a different orientation. Representative distances between Fe-DHB and the TBDT substrate metal ions are shown, colored as in A.

### The amino acids at the Fiu external binding site are important for iron acquisition *in vivo*

To determine to role of the amino acids that define the putative Fiu external substrate binding site, we assessed the functionality of variants of Fiu with mutations in this region (Figure 5, Figure S6). Small side-chains around the cavity (contributed through alanine, serine and threonine) were mutated to the bulky tryptophan to sterically occlude the binding pocket, while larger side-chains defining the pocket were mutated to alanine. Plasmid-borne constructs of these mutant Fiu proteins were transformed into the *E. coli* BW25113 strain Δ6 (Figure 1B,C). To test for restoration of iron transport activity, the transformed strains were streaked onto LB agar with 2’2-bipryidine and scored for growth (Figure 5, Figure S6). Mutation of phenylalanine 105 (F105A), glutamate 108 (E108A) and arginine 142 (R142A) grossly affected the function of Fiu (Figure 5, Figure S6). Fiu(E108A) was non-functional, displaying growth identical to the negative control. Fiu(F105A) and Fiu(R142A) displayed minimal complementation (Figure 5, Figure S3). Two other mutations, threonine 113 to tryptophan (T113W), and serine 139 to tryptophan (S139W), also exhibited some defect in function compared to wildtype Fiu (Figure 5, Figure S3).

**Figure 5.**
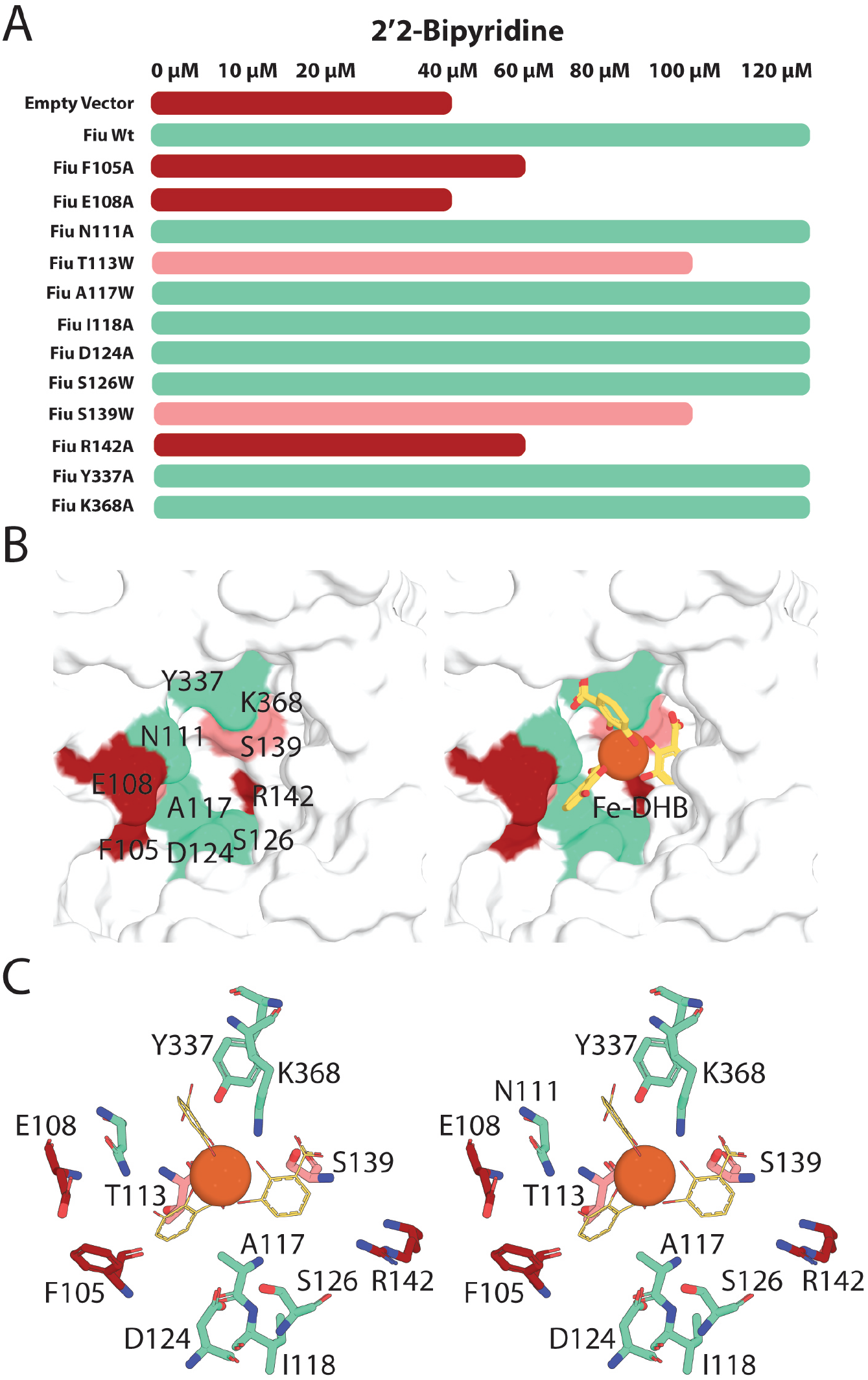
The effect of mutations to the Fiu putative external Fe-DHB binding site on the function of Fiu *in vivo*. (A) The effect of Fiu binding site mutations of the ability of pBAD24Fiu to complement *E. coli* BW25113 Δ6, grown on LB agar with increasing concentrations of 2,2’-Bipyridine. (B) The location of mutations from panel A in the external binding site of Fiu, shown as a surface model and color coded for effect of Fiu function. Amino acids and labelled in the left panel, Fe-DHB is present in the right panel (C) A stereo image showing the residues mutated to the Fiu external binding site in a stick representation, colored as panel A. The Fe-DHB complex is shown as a line and sphere representation for DHB and Fe^3+^ respectively.

Residues F105 and E108 are located on plug domain loop 1 within the predicted Fe-DHB binding pocket (Figure 5C). F105 is ordered in both crystal states and forms a hydrophobic pocket, which shelters the aromatic carbons of one of the DHB monomers of the Fe-DHB complex in our docked structure. E108 is disordered in the open state of Fiu and is not within bonding distance of docked Fe-DHB (5.3 Å), but may form interactions with polar substrate groups *in vivo*, or generally stabilize the binding pocket (Figure 5B, C). R142 is minimally surface exposed and unlikely to interact directly with catecholate substrates (Figure 5B). However, it is located in plug domain loop 2 and forms hydrogen bonds with the carbonyl groups of amino acids D136 and G138. These interactions stabilize this loop, which defines the lower section of the substrate binding pocket and this may account for the phenotypic effect of the mutation. Mutation of N111, Y337 and K368 that directly interact with Fe-DHB in our docked structure did not affect Fiu function in the in vivo assay (Figure 5). This may reflect a lack of precision in our docked model, or the fact that single point mutations of substrate interacting residues may be insufficient to effect substrate import, as has been reported for other TBDTs [41]. Given several single mutations in this region dramatically effect Fiu function, these data provide evidence that the external binding site identified in our docking is important for Fiu function.

## Discussion

*E. coli* BW25115 possesses three outer membrane TBDTs known to be responsible for the transport of Fe-catecholate complexes, Fiu, FepA and Cir [32, 51]. In this work, we show that despite sharing substrates related by a catecholate functional group, Fiu is only distantly related to FepA and Cir. This demonstrates that TBDTs from different linages have the ability to transport similar substrates, and they may have arrived at this specificity via convergent evolution.

FepA is a high affinity transporter for enterobactin, a siderophore endogenously produced by *E. coli* [31, 36]. While FepA transports enterobactin, like Fiu and CirA it is also able mediate the import of iron in complex with monomeric catecholates, albeit with a longer affinity [32]. As Fiu does not transport enterobactin, it is unclear whether its major physiological role is the transport of iron in complex with monomeric catecholate molecules or in the import of a hitherto unidentified xenosiderophore. In this work we show that Fiu supports the growth of *E. coli* in the absence of an exogenously supplied substrate, likely through the import of Fe-catecholate compounds generated through breakdown of enterobactin. However, the ability of Fiu to transport iron under these conditions is inferior to that of FepA and does not support growth under stringent iron limitation. The relatively poor ability of Fiu to transport iron under these conditions suggests transport of enterobactin breakdown products is a secondary function for this transporter, leaving its high affinity substrate to be identified. This hypothesis is further supported by our inability obtain a bonafide complex between Fiu and either Fe-DHB or Fe-DHBS, despite identifying conditions that were clearly compatible with substrate crystallization.

The crystal structure and associated analysis that we present further reinforces the differences between Fiu and FepA. FepA and its closely related homologue PfeA from *P. aeruginosa*, initially bind enterobactin an external binding site formed by extracellular loops that entirely enclose the entrance of these transporters β-barrel [42, 50]. Fiu lacks the extracellular loop structure required for this external binding site and our docking analysis suggests it binds its substrate deep in its internal cavity, at a location shared among other TBDTs [38, 41–49].

Structural analysis of Fiu reveals that a number of extracellular loops are selectively ordered and this selective order is shared among the homologous receptors PiuA and PiuD [30, 33]. The observed crystal states may represent conformational transitions that the receptor undergoes during substrate binding and transport. Specifically, the external plug loop of Fiu in our structures selectively gates a large cavity in the lumen of the transporter. Our docking shows that this cavity is large enough to accommodate a Fe-catecholate siderophore complex. Additionally, based on predictions of the liable subdomain of the N-terminal plug [37, 39], this cavity is in the path of substrate transport (Figure 2C). This selectively ordered Fiu plug loop substrate may allow its substrate to enter the internal cavity prior to the removal of the liable subdomain. If the external plug loop then transitioned to an ordered state during subdomain removal by TonB, it would prevent channel formation through Fiu during import, and prevent the non-specific import of deleterious substances. Despite their significant structural differences and distinct initial binding sites, the essence of this mechanism may be shared with FepA, which possesses a second enclosed binding site deeper inside the barrel of the transporter [42, 50].

Gram-negative bacteria can exhibit a formidable resistance to antibiotics, largely due to the protective semi-permeability of the outer-membrane [2, 52, 53]. Catecholate containing antibiotics use molecular mimicry to facilitate their import into bacteria via Fiu and related transporters [25]. As these transporters present in diverse bacterial groups, siderophore mimicking antibiotics are attractive lead compounds for the development of new therapeutics to treat Gram-negative bacterial infections, representing an area of considerable recent interest for antibiotic development [28, 29, 54]. In *Acinetobacter baumannii* and *Pseudomonas aeruginosa*, the Fiu homologue PiuA has been shown to greatly promote the susceptibility to the 3,4-dihydroxypyridine containing sulfactam BAL30072 [30, 55].

By investigating the structural basis for substrate binding and import by Fiu, this work will assist in the exploitation of Fiu and related transporters, as a conduit for catecholate-like antibiotics into the bacterial cell.

## Experimental Procedures

### TBDT phylogeny construction, Fiu homologue search and clustering analysis

To determine the phylogenetic relationship between Fiu and TBDTs of know structure and/or function, a subset of TBDT sequences were selected and sequences were obtained from the NCBI database. Sequences were aligned using the Clustal algorithm and the alignment was utilised to build a bootstrapped phylogenetic tree (100 repetitions), which was visualized using the FigTree software [56].

A HMMER search using the Fiu sequence from *E. coli* BW25113 as the search query was established using the stable and unbiased proteome dataset RP55 [57, 58]. The search was restricted to sequences with an E-value of less than 1e-75, thereby limiting the outcome to 502 sequences (SI Index Table S3). For sequence clustering, classification used an all-against-all BLAST clustered based on pairwise similarities and visualized with CLANS [59] with a E value cut off of 1×10^-120^. To assess the similarly of sequences identified this search to other TBDTs present in *E. coli* BW25113 sequences for FhuA, FecA, CirA, FepA, BtuB, YddB and YncD were added to this dataset and clustering was performed with an E value cut off of 1×10^-120^.

### Construction of multiple TBDT knockout *E. coli* BW25113 strain

*E. coli* BW25113 mutants were created using the λ-red system [60]. Kanamycin resistance cassettes flanked by 300bp of genomic DNA corresponding to regions in each of the genes encoding TBDTs of interest were amplified using specific mutants from the *E. coli* mutant Keio collection [61] as templates. Primers are summarized in Table S5.

The host strain *E. coli* BW25113 was transformed with the λ-red recombinase plasmid pKD46 [60], grown at 30 °C (LB broth + 100 μg.ml^-1^ ampicillin) to an OD 600nm of 0.1 before λ recombinase was induced by the addition of 0.2 % L-arabinose. Cultures were thereafter grown at 30 °C until OD 600nm 0.6-0.8 and transformed using the room temperature electroporation method [62]. Briefly, bacterial cells were isolated by centrifugation at 3000 g for 3 minutes, and washed twice with a volume of sterile 10 % glycerol equal to the volume of the culture used. Cells were then resuspended in 10 % glycerol to a volume of 1:15 of the volume of culture. 100-500 ng of PCR amplified Kan^R^ KO cassette for the gene of interest was then added to 100 μl of the resuspended bacteria and the mixture was electroporated. 1ml of LB broth was added to the cells post electroporation, and the culture was recovered at 37 °C for 1 hour, before plating onto LB agar + 30 μg.ml^-1^ Kanamycin. PCR was used to validate that colonies did indeed have the Kan^R^ cassette in place of the gene of interest.

To remove the Kan^R^ cassette, deletion mutant strains were transformed with the plasmid pCP20 [63] containing the “flippase cassette”. Cells were growth under either ampicillin (100 μg.ml^-1^) or chloramphenicol (30 μg.ml^-1^) selection to maintain the plasmid. For removal of the Kan^R^ cassette, a single colony of the mutant strain was used to inoculate 1 ml of LB broth (no selection). The culture was growth overnight at 43 °C to activate expression of the flippase gene. This culture was then subject to 10-fold serial dilution in sterile LB and plated onto LB agar with no selection. Resulting colonies were patched onto LB agar containing kanamycin, chloramphenicol or no selection. PCR was used to validate that colonies unable to grow of kanamycin or chloramphenicol, but growing in the absence of selection, did indeed represent successful removal of the Kan^R^ cassette. This process was repeated sequentially to derive strains multiply defective in up to six TBDT receptors. The order of deletion was ΔFhuA, ΔFecA, ΔCirA, ΔFepA, ΔFhuE and then ΔFiu. Mutant strains created in the process were designated Δ1, Δ2, Δ3, Δ4, Δ5 and Δ6, based on the number of receptors deleted.

### Testing the growth of *E. coli* BW25113 TBDT deletion strains under iron limiting conditions

*E. coli* BW25113 deletion strains (Δ4, Δ5 and Δ6) grew poorly on LB agar. To ameliorate this phenotype all mutant strains were maintained on LB agar + 250 μM Fe(II)SO_4_. All deletion strains grew well under these conditions. To test the ability of mutant strains to grow under iron limiting conditions, strains were grown in LB broth until stationary phase. Cells were harvested from 0.5 ml of this stationary phase culture and the supernatant was removed. Cells were resuspended in 0.5 ml of 1x M9 salts and a minimal quantity of this suspension was streaked onto LB agar containing 0-150 μM 2’2-bipyridine. Plates were incubated at 37 °C overnight and growth was observed and scored.

### Complementation of TDBT receptor mutants with wildtype and mutant Fiu

The open reading frame for Fiu, including the sequence encoding the signal peptide, was amplified from *E. coli* BW25113 by PCR (Table S5) and cloned into the pBAD24 plasmid, at EcoRI and HindIII restriction sites. The resulting vector, designated pBAD24Fiu, was then transformed into *E. coli* BW25113 Δ6 and maintained using 100 μg.ml^-1^ ampicillin. In order to test for complementation, *E. coli* BW25113 TBDTΔ6 pBADF24iu was streaked onto LB agar 0.2 % arabinose, 100 μg.ml^-1^ ampicillin and 0-120 μM 2,2’-Bipryidine. Growth under these conditions was compared to that of *E. coli* BW25113 Δ6 containing pBAD24 as a vector control. Mutants of the putative Fiu substrate binding site were created via whole plasmid mutagenesis using pBADFiuCom as the starting vector [64]. The mutations were introduced using the primer sequences provided in Table S6. The sequence of the pBAD24Fiu template and introduction of the specified mutations in the resultant plasmids were confirmed by Sanger sequencing. Mutant plasmids were transformed into *E. coli* BW25113 TBDTΔ6 and maintained tested for function as described above for pBAD24Fiu.

### Protein Expression and Purification

DNA encoding the mature form of Fiu lacking the signal peptide was amplified from *E. coli* BW25113 using primers shown in SI Index Table S6. NcoI and XhoI restriction sites incorporated into the primers were used to clone the DNA fragment into a modified pET20b vector with a 10x N-terminal His-tag followed by a TEV cleavage site. The resulting plasmid was transformed into *E. coli* BL21 (DE3) C41 cells, and protein expression was induced in cultures in terrific broth (12 g tryptone, 24 g yeast extract, 61.3 g K_2_HPO_4_, 11.55 g KH_2_PO_4_, 10 g glycerol) with 100 mg.ml^-1^ ampicillin for selection. Cultures were grown at 37 °C until OD_600_ of 1.0, induced with 0.3 mM IPTG and growth for a further 14 hours at 25 °C. Cells were harvested by centrifugation, lysed with a cell disruptor (Emulseflex) in Lysis buffer (50 mM Tris, 200 mM NaCl, 2 mM MgCl_2_) plus 0.1 mg.ml^-1^ Lysozyme, 0.05 mg.ml^-1^ DNAse1 and Complete protease cocktail inhibitor tablets (Roche). The resulting lysate was clarified by centrifugation at 20,000 g for 10 minutes, the supernatant was then centrifuged for a further 1 hour at 160,000 g to isolate membranes. The resultant supernatant was decanted and the membrane pellet was suspended in Lysis buffer, using a tight fitting homogeniser. Once homogenized, the membrane fraction was solubilised by the addition of 10 % Elugent (Santa Cruz Biotechnology) and incubated with gentle stirring at room temperature for 20 minutes. Solubilised membrane proteins were clarified by centrifugation at 20,000 g for 10 minutes. The supernatant was applied to Ni-agarose resin equilibrated in Ni binding buffer (50 mM Tris, 500 mM NaCl, 20 mM Imidazole, 0.03% Dodecylmatoside (DDM) [pH7.9]). The resin was washed with 10-20 column volumes of Ni binding buffer before elution of the protein with a step gradient of, 10, 25 and 50, 100 % Ni gradient buffer (50 mM Tris, 500 mM NaCl, 1 M Imidazole, 0.03 % DDM [pH7.9]). Fiu eluted at the 50% and 100 % gradient steps. Eluted fractions containing Fiu were pooled and applied to a 26/600 S200 Superdex size exclusion column equilibrated in SEC buffer (50 mM Tris, 200 mM NaCl, 0.03 % DDM [pH 7.9]). To exchange Fiu into Octyl β-D-glucopyranoside for crystallographic analysis fractions containing Fiu were pooled an applied to Ni-agarose resin, equilibrated in βOG buffer (50 mM Tris, 200 mM NaCl, 0.8 % Octyl β-D-glucopyranoside [pH 7.9]). The resin was washed with 10 column volumes of β-OG buffer before elution with βOG buffer + 250 mM imidazole. Fractions containing Fiu were pooled and 1 mg.ml^-1^ 6 x histidine tagged TEV protease and 1 mM DTT were added. This solution was then dialysed against of β-OG buffer at 4-6h at 20 °C to allow TEV cleavage of the 10 x histidine tag and removal of excess imidazole. The solution was then applied to Ni-agarose resin, to remove TEV protease and the cleaved histidine peptide. The flow through containing Fiu from this step was collected concentrated to 14 mg/ml^-1^ in a 30 kDa cut-off centrifugal concentrator and snap frozen and stored at −80 °C.

### Protein Crystallisation, Data Collection and Structure Solution

Purified Fiu in β-OG buffer was screened using commercially available crystallisation screens (approximately 600 conditions). Hexagonal crystals formed in the JCSG screen in 1 M LiCl, 20 % PEG 6000 and 0.1 M Tri-sodium citrate pH 4.0 [65]. These crystals diffracted poorly (>3.3 Å) and suffered from considerable anisotropy. In order to improve diffraction, Fiu in the above condition was subjected to an additive screen (Hampton Research). Hexagonal crystals grew with many additives, however in the presence of 5% polypropylene glycol P400 (PPG 400) flat diamond shaped plates formed. Crystals diffracting to 2.1 Å in the space group C222_1_ were obtained and, despite relatively low sequence identity (32%) between Fiu and the catecholate receptor PiuA from *Acinetobacter baumannii* (5FP1), a molecular replacement solution was obtained using phaser with the crystal structure of the PiuA receptor as a starting model [30, 66]. The model was built and refined using the phenix package and coot [67, 68]. The majority of the Fiu polypeptide chain could be modelled into the available density, however loops 7-9 and the plug domain loop were disordered in this structure. The Fiu model from these crystals was designated crystal state 1.

In order obtain Fiu is additional crystal states, purified Fiu in β-OG buffer with 5% PPG 400 was rescreened for crystallisation (∼600 conditions). Fiu crystallized in multiple addition conditions in the presence PPG400. Using crystals derived from these screens the structure of Fiu was solved in 2 further crystal forms: *P1* form at 2.9 Å in 20 % PEG 3350, 0.2 M Na_2_ Malonate, 0.1 M Bis-Tris Propane [8.5 pH] and C2_1_ form at 2.5 Å in 0.1 M Tris, 20 % PEG 6000, 0.2 M NaCl [pH 8.0] (+4 % PPG400). In these crystal forms all loops of Fiu were ordered and conformationally analogous, allowing complete tracing of the Fiu amino acid sequence. Fiu modelled from these crystals was designated crystal state 2.

Co-crystallisation between Fiu and Fe-DHB or Fe-DHBS was performed by adding 300 μM – 1mM to purified Fiu in β-OG buffer +/-5 % PPG 400 to at a final concentration of ∼100 μM (8mg/ml). Crystal screening was performed as above. Crystals were harvested directly from screening trays, mother liquor was removed by wicking and crystals were cryocooled in liquid N_2_ at 100 K. For soaking experiments, Fiu crystals from crystal states 1 and 2 were transferred to well solution containing 1 mM Fe-DHB and Fe-DHBS incubated in this solution for between 1-5 minutes cryocooling.

### *In silico* docking of Fiu substrates with the Fiu crystal structure

In order to determine potential ligand bindings sites in the Fiu crystal structure, an *in silico* docking approach was applied using Autodock Vina, within the Chimera software package [40, 69]. Coordinates for the iron in complex with three molecules of 2,3-Dihydroxybenzoic acid (Fe-DHB) was obtained from 3U0D: The structure of human Siderocalin bound to the bacterial siderophore 2,3-DHBA. Coordinates for BAL30072 and Pirazmonam were obtained from the PubChem database (https://pubchem.ncbi.nlm.nih.gov/). Coordinates for Fiu were taken from molecule A of the C2_1_ crystal form for theclosed state and from molecule A of the C2221 crystal form for the open state. Ligand and Fiu coordinates were imported into Chimera and optimised using the Dock Prep utility. Docking was performed using the Autodock Vina dialogue, two box sizes were utilised for docking (Box 1 = 46.5, 56.0, 45.5 Å, Box 2 = 46.5, 35.2, 45.5 Å). Box 1 encompassing the entire extracellular portion of Fiu and all cavities accessible from the extracellular environment. Box 2 excluded the large cavity in Fiu, gated by the extracellular plug domain. A total of 10 binding modes were sought for each docking run, with a search exhaustiveness of between 8 and 300 and a maximum energy difference of 3 kcal/mol. Docking solutions did not differ significantly as a result of changes to search exhaustiveness. Docking solutions were visually inspected and the highest rated solution was used for main figures and for discussions.

### Analysis of TBDT ligand binding sites

The Protein Data Bank (PDB) was manually searched for structural coordinates of TBDT in complex with substrate compounds TBDT receptor complexes were aligned to the crystal structure of Fiu (Table S5), based on the TBDT chain using the ‘super’ command in pymol. The location bound substrates was determined by manual inspection in Pymol. The location and volume of Fiu substrate binding cavities was estimated using CASTp [70].

### Data availability

All crystallographic coordinates and associated structure factors produced in this study are available in the Protein Data Bank (PDB) with the accession codes: 6BPM = Fiu in space group C21, 6BPN = Fiu in space group C2221, 6BPO = Fiu in space group P1

## Acknowledgments

This research was undertaken in part using the MX2 beam-line at the Australian Synchrotron, part of ANSTO, and made use of the Australian Cancer Research Foundation (ACRF) detector. This research was undertaken on the MX1 and MX2 beamlines at the Australian Synchrotron, part of ANSTO (CAP12312). We would like to thank the Monash Crystallisation Facility for their assistance with sample characterisation, crystallographic screening and optimization. We would like to thank Dr Laura C. McCaughey and Dr Inokentijs Josts for their critical reading of the manuscript and resulting suggestions. The work was funded by the Australian Research Council (ARC; FL130100038) and the National Health & Medical Research Council (NHMRC Program in Cellular Microbiology, 1092262). R.G. was funded by a Sir Henry Wellcome Fellowship award (106077/Z/14/Z). T.L. is an ARC Australian Laureate Fellow (FL130100038).

## Author contributions

Conceived and designed the experiments: RG, TL

Performed the experiments: RG

Analyzed the data: RG, TL

Contributed reagents/materials/analysis tools: RG, TL

Wrote the paper: RG, TL

## Supplemental Data

**Table S1 List of sequences homologous to Fiu identified in HMMER search.**

**Table S2 Sequence identity matrix of Fiu homologues from Enterobacteriaceae.**

**Table S3 Data collection and refinement statistics for Fiu crystal structures.**

**Table S4 Fiu Fe-DHB Autodock docking statistics**

**Table S5 The crystal structures of TBDT-substrate complexes utilised for binding site analysis.**

**Table S6 Oligo nucleotide primers used in this study**

**Table S7 Strains and plasmids used or generated in this study**

**Figure S1.**
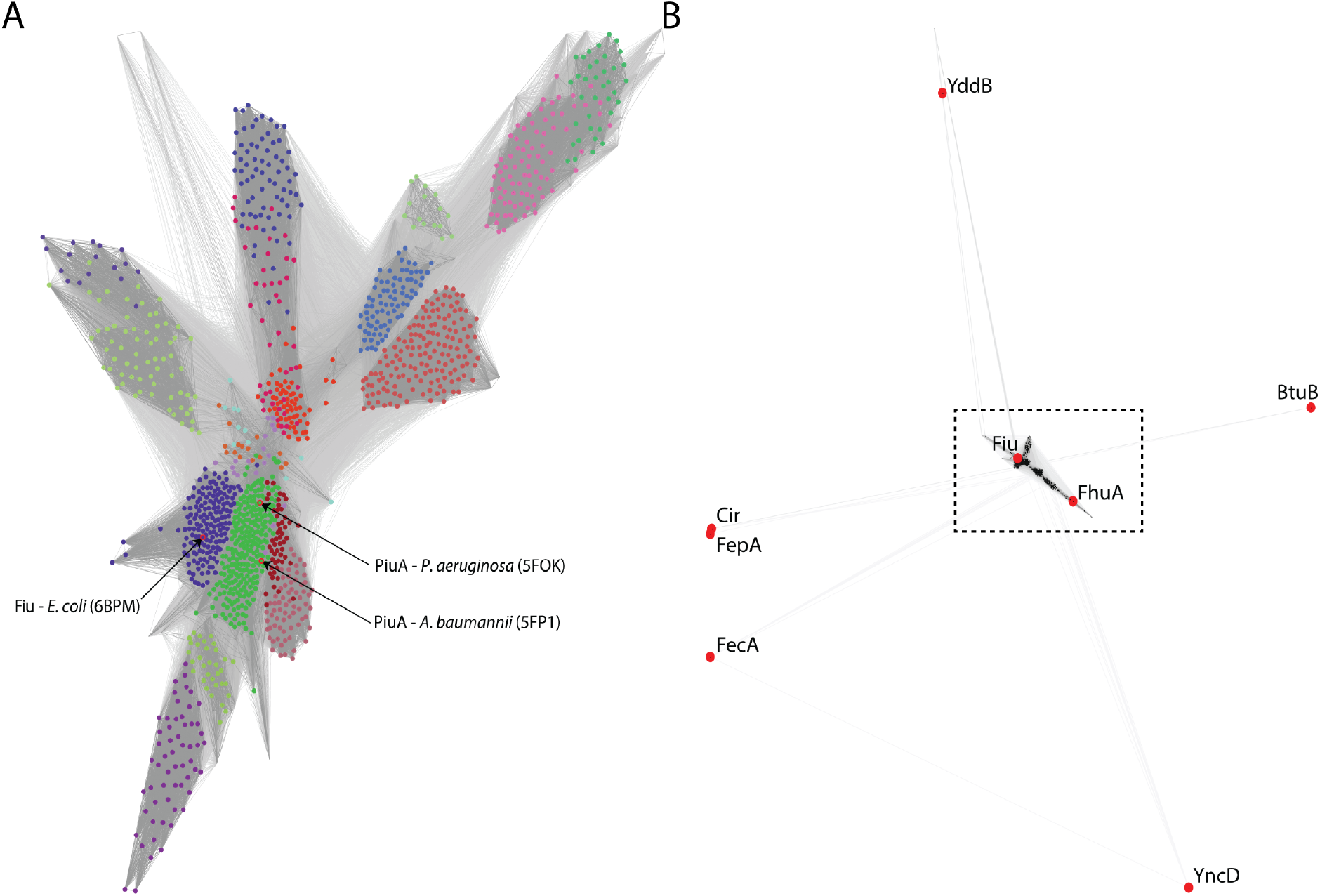
CLANS clustering of Fiu homologues identified by HMMER search and other TBDTs from *E. coli*. (A) CLANS clustering analysis [59] of Fiu homologue sequences identified by HMMER search (Table S1), location of point corresponding to Fiu and PiuA are indicated. Sequences in (B) are spiked with TBDT sequences from *E. coli* BW25113 to show relationship between these receptors and the Fiu homologues identified in the search.

**Figure S2.**
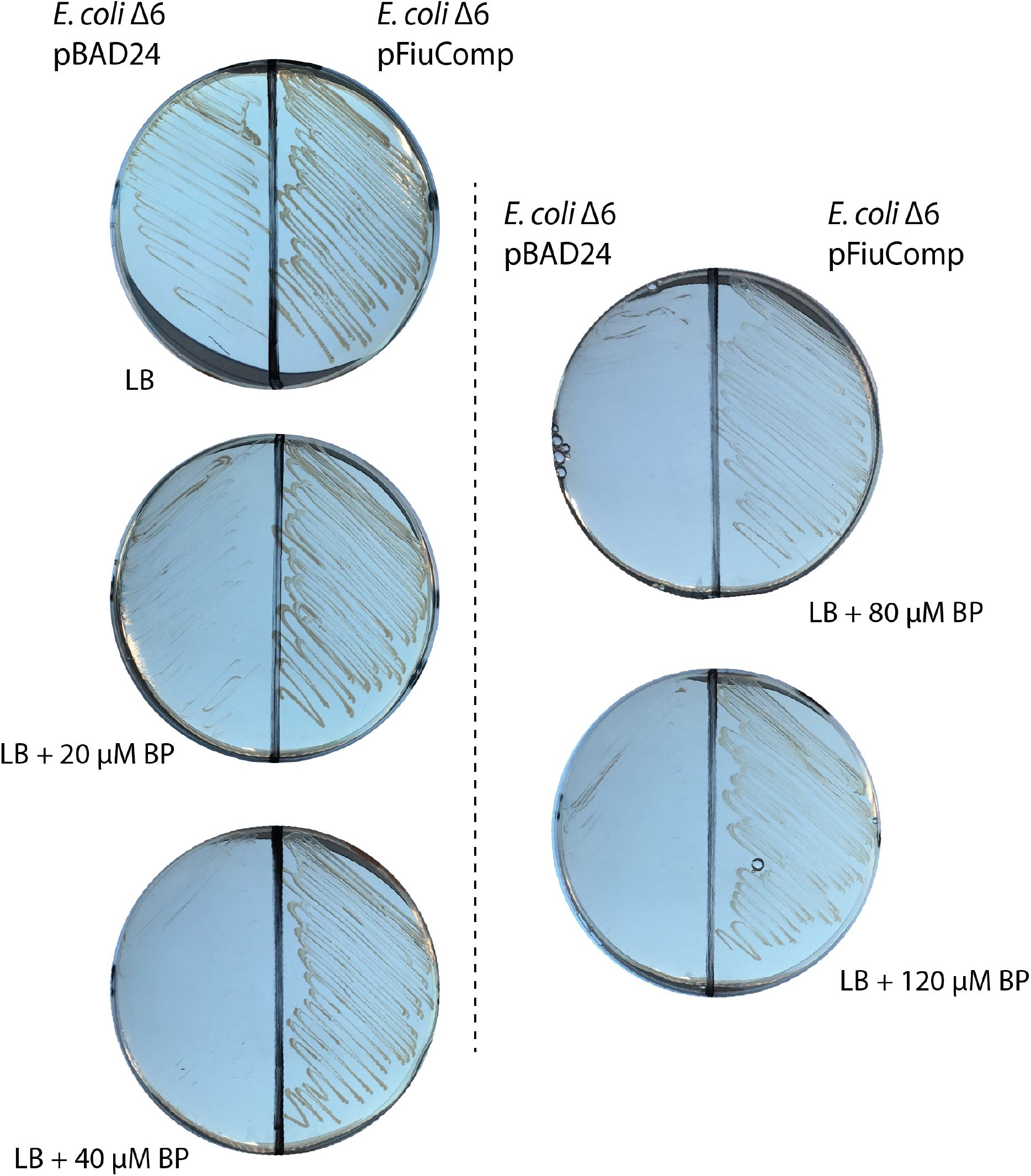
Complementation of *E. coli* BW25113 TBDTΔ6 with plasmid derived Fiu. Growth of *E. coli* BW25113 TBDTΔ6 with complementation vector (pBAD24Fiu) or vector control (pBAD24) on LB agar with 0-120 μM 2,2’-Bipryidine (BP). Vector control is unable to grow at a concentration of BP > 20 μM, while complemented strain can grow at a concentration of BP < 120 μM.

**Figure S3.**
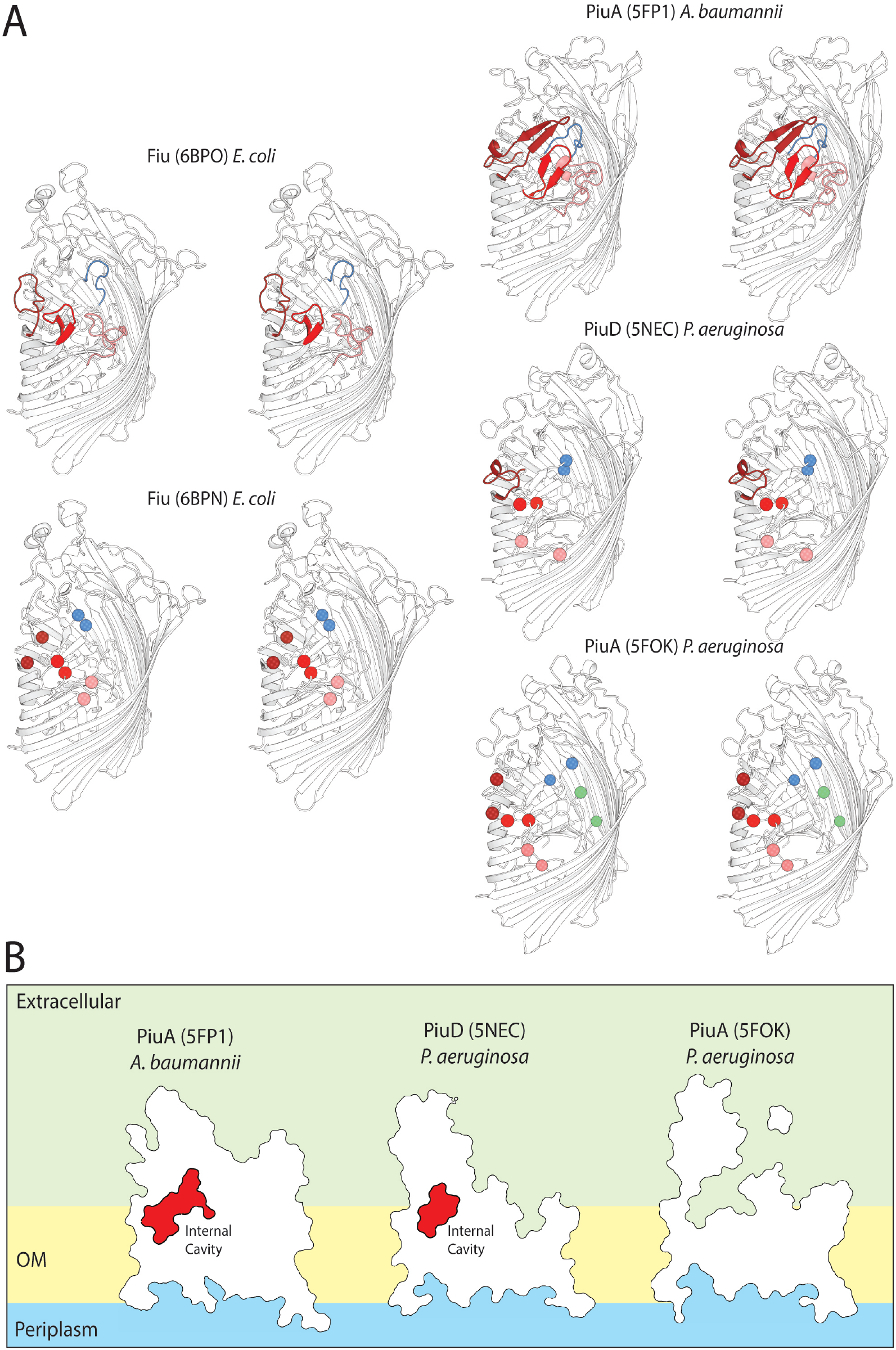
Extracellular loop stability in crystal states of Fiu and PiuA/PiuD. (A) A stereo cartoon view illustrating the variably ordered extracellular barrel and plug domain loops in Fiu crystal states 1 and 2 and PiuA and PiuD crystal structures from *P. aeruginosa* and *A. baumannii*. The crystal structures of PiuA and Dare analogous to the variable loop order/disorder observed for Fiu. The plug loop is shown in blue, extracellular loops 7, 8 and 9 are shown in pink, red and brick red respectively (termini shown as spheres where the loop is disordered) (B) A cutaway outline representation of PiuA and PiuD, showing the presence of a selectively gated internal cavity.

**Figure S4.**
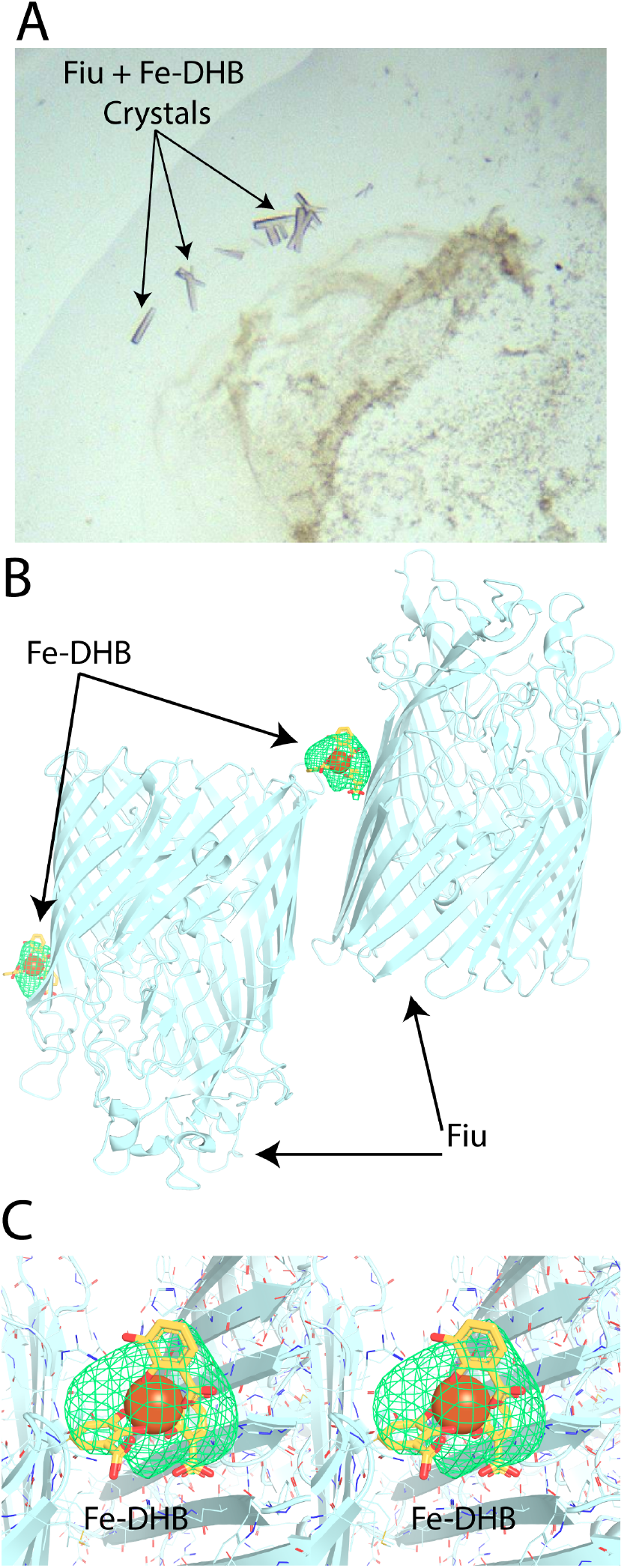
Co-crystallisation between Fiu and Fe-DHB complex. (A) Crystals of Fiu obtained in the presence of Fe-DHB, with violet color characteristic of a Fe-Catecholate complex. (B) The resultant asymmetric unit from these poorly diffracting crystals (anisotropic 3.2-5 Å), containing 2 molecules pf Fiu in complex with 2 Fe-DHB complexes. Fiu is shown in a cartoon representation, while Fe-DHB is shown as a stick/sphere model modelled into Fo-Fc density contoured to 3 σ. (C) A zoomed view of the central Fe-DHB complex from panel B, with Fiu sidechains shown as lines. Fe-DHB is coordinated by a patch of positively charged residues on one Fiu molecule.

**Figure S5.**
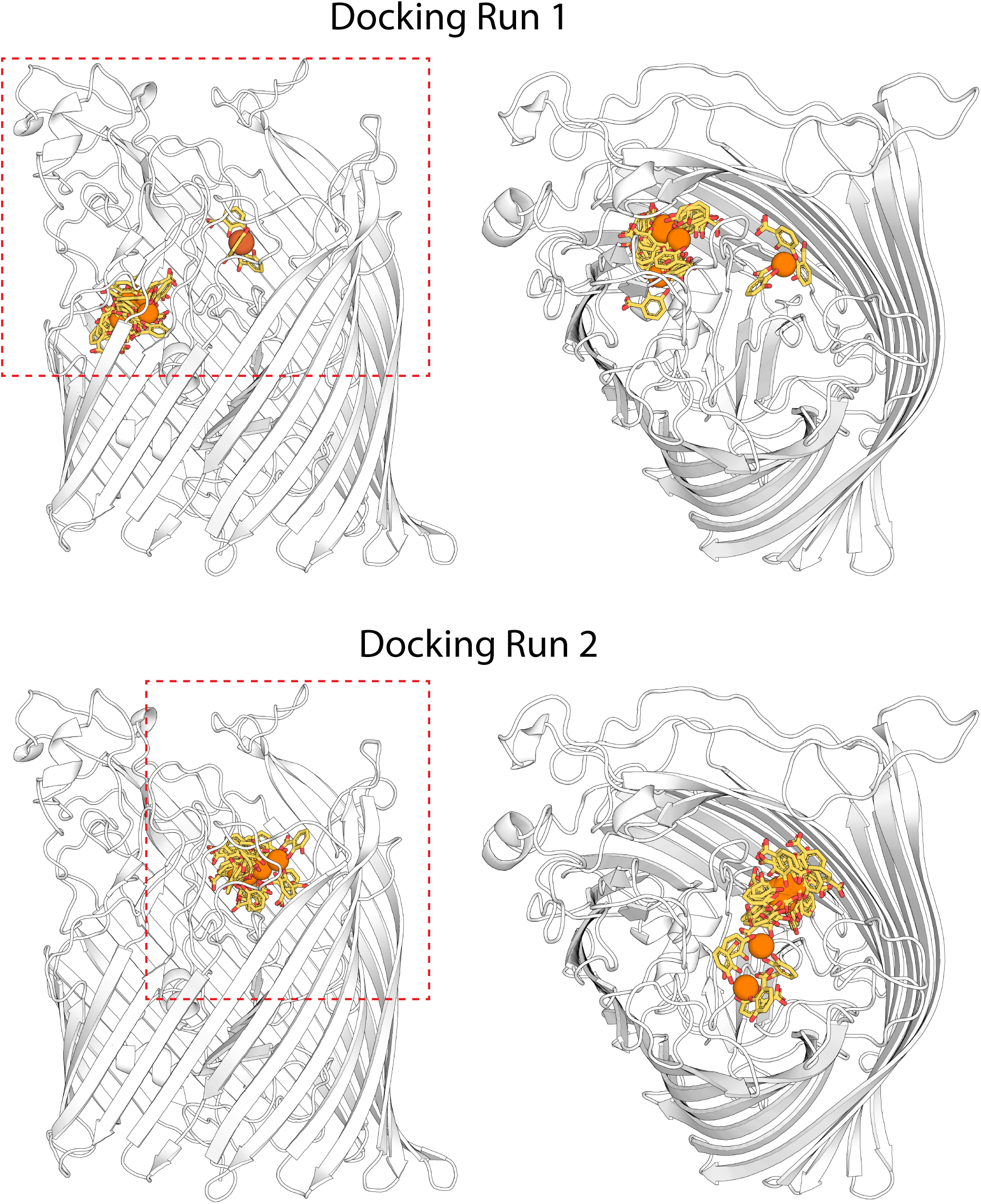
All docking modes obtain between Fiu and Fe-DHB. All 9 docking modes obtained for docking runs 1 and 2, shown with Fe-DHB as a stick/sphere model with Fiu with a cartoon representation. The dashed red box shows docking search area along the Z-axis relative to the membrane. In both docking runs the majority of binding modes, including the top ranked mode cluster at a single location.

**Figure S6.**
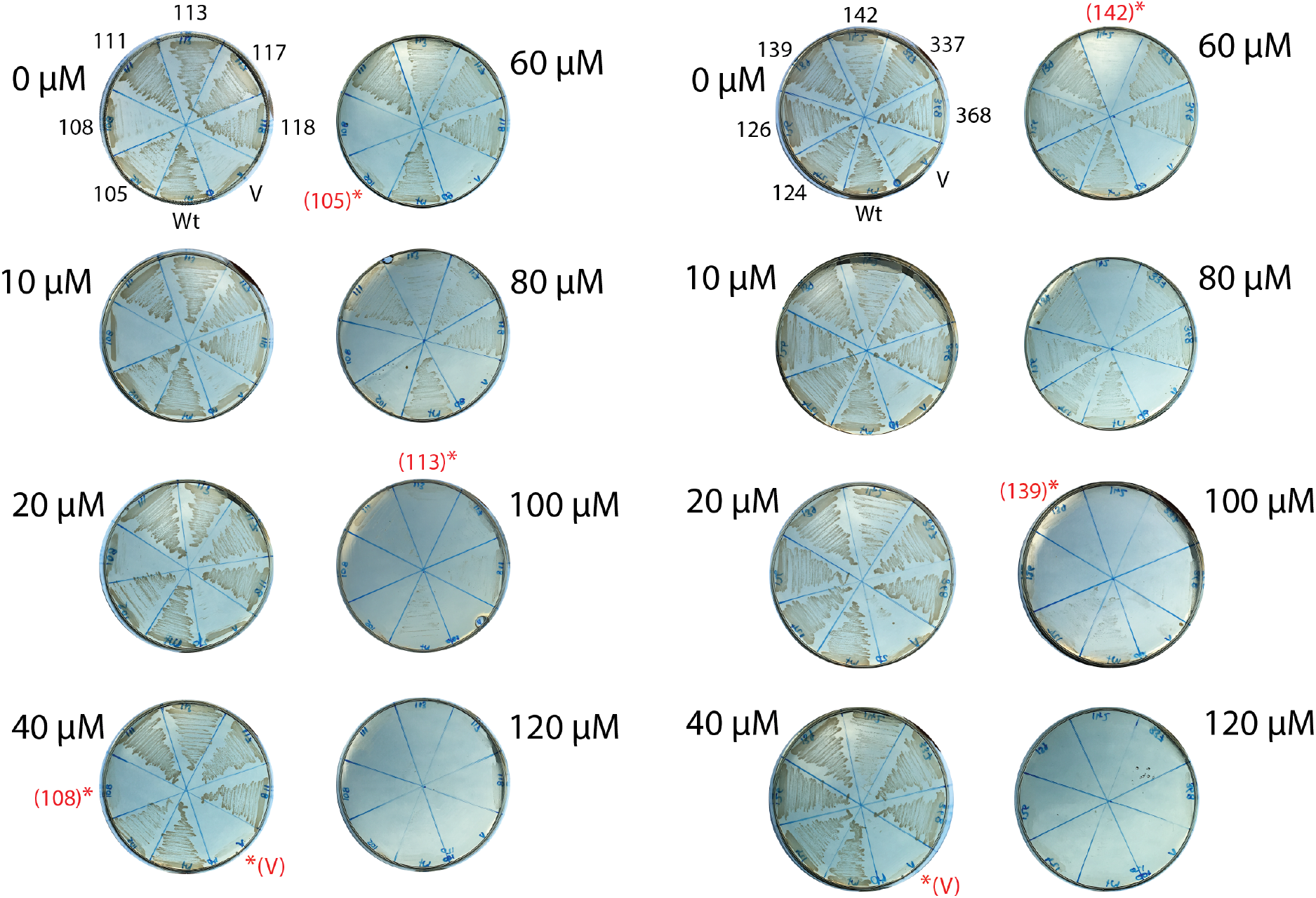
Effect of substrate binding site mutations on *in vivo* function of Fiu. The growth of *E. coli* BW25113, containing complementation vector (Wt), vector control (V) or complementation vector mutants (numbers) on LB agar with 0-120 μM 2’2-bipryidine (BP). Mutant numbers represent position of the amino acids mutated (F105A, E108A, N111A, T113W, A117W, Y118A, D124A, S126W, S139W, R142A, Y337, L368A). Starred sections show the maximum BP concentration at which the corresponding mutated construct or control is able to grow.

**Structural Coordinates for Fiu docked with Fe-DHB (provided with final submission).**

